# The strong grip of childhood conditions in older Europeans

**DOI:** 10.1101/267526

**Authors:** Gindo Tampubolon, Maria Fajarini

## Abstract

Among older Europeans grip strength has been found to be marked by a disadvantaged adulthood. Across the Channel, among older Britons gait speed as another measure of physical function has been found to be marked by disadvantaged childhood. Using the Survey of Health, Ageing, and Retirement in Europe (2004-2013), we studied whether childhood poverty led to Europeans aged 50 to 104 years having a weaker grip. We then drew their trajectories of repeatedly measured grip strength to discern a steeper decline among the childhood poor. Retrospective childhood poverty some four to nine decades in the past was treated as a latent construct following the above literature; attrition during repeated measurements is handled using inverse proportional to attrition weighting. The data showed the childhood poor to have a weaker grip for half a century in later life. However, they do not show a steeper decline. Most important, by contributing to levels of grip strength in later life, adult condition holds the potential to shape the strong and long arm of childhood condition. The results are another impetus to eliminate childhood poverty to ensure healthy ageing Europeans.

## Introduction

Physical functioning is a key driver for the wellbeing of older people, with grip strength and gait speed its two important and well-characterised markers [1]. Therefore, maintaining high levels of physical function throughout later life is singled out as an objective for public health in response to the ageing population challenge [2]. It is also important because with life expectancy at 60 years extending secularly, the welfare implication of impaired physical function in later life is considerable. Long term care of physical disabilities in later life is costly. In the Netherlands and Sweden in 2011, it costs more than 3.5% of their gross domestic products [2]. We therefore aimed to draw trajectories of physical function of Europeans aged 50 to 104 over an extended period.

Grip strength has been repeatedly shown to predict incident disability, morbidity, and mortality [3, 4]. Thus a recent study explored its predictors among Europeans aged 65 to 90 using the Survey of Health, Ageing, and Retirement in Europe (SHARE, 2004-2013), examining the roles of parental occupation and individual occupation at midlife [5]. The authors found that grip weakens linearly with age with steeper slopes among men than women; similar results had been found earlier in a cross-section of Europeans aged 50 and older [6] and in Danes aged 46 to 102 [7]. In addition, men with elementary or lower occupation at midlife had a weaker grip at ages beyond 65, though there was no evidence that they experienced a steeper decline. Beyond age and sex variations, an earlier study of SHARE found that grip strength varied considerably across individual height and geographic region (north – south), advising that these variables should be adjusted for [6].

Another report used gait speed as a measure of physical function in its sister study, the English Longitudinal Study of Ageing (ELSA), appraising the role of an even earlier stage in life course: childhood condition [8]. The author found that material poverty during childhood associates with slower gait in Britons aged 50 to 90 years, with material poverty indicated by lack of essential facilities, overcrowding, and number of books in the childhood home, as well as financial hardship during childhood. Notably, childhood information was elicited retrospectively, collecting potentially inaccurate information [9], and requiring new methods based on latent construct to deal with inaccurate information [8]. Examined with the new methods, the data showed that childhood poverty was associated with lower levels of health status in later life overall: slower gait, poorer memory, and more depression. The mechanism invoked to link the childhood condition and later life emphasised the broader effects early life adversity can have. The results evinced the long arm of childhood condition across the spectrum of health from physical to mental health.

To understand more about the long arm of childhood condition [10], four improvements can be made. First, most empirical studies are content with explaining levels of health, effectively associating a stage in childhood and a time in later life. The study of older Britons above for example explained the levels of gait speed, episodic memory, and depression by childhood condition. No attempts was made at explaining their rates of change. But surely it is more fruitful to understand whether childhood poverty puts people onto a trajectory of steeper decline. So far, the limited evidence shows no steeper decline among those with disadvantaged childhood or midlife [5].

Second, with some exceptions [5, 8], most empirical studies stopped at adulthood. No doubt, this is a function of available data. Although theory suggests that a disadvantaged childhood can be compensated for in adulthood and midlife such that later life health is freed from childhood condition [11], very little evidence is furnished about older people and their childhood. On the other hand, epigenetic change in early life is posited to have a stable effect well into later life [12]. Once biological imprinting has transpired through DNA methylation and histone modification, the effect of childhood condition can persist. Therefore, more empirical investigation is necessary to examine whether childhood condition reach into health trajectories in later life.

Third, information about childhood condition of the oldest old [13] is rarely available in prospective survey. This lack is felt more strongly if a nationally representative sample is required. ELSA collected rich information about people aged 50 years and over prospectively, except when it comes to information about their childhood which was collected retrospectively. This information may not be entirely accurate. For instance, among 50 year old Britons (who had been prospectively followed since birth), when asked about the numbers of people and bedrooms in their childhood home, only one in three got both right [9]. Fortunately, new methods to work with such inaccurate retrospective information have been proposed and subsequently shown to work with these kinds of data; such methods need to be applied more often [8].

Lastly, studying older people over time to draw their trajectories of physical function inevitably faces attrition problems since older people tend to attrite from a longitudinal study due to worse health function [14, 15]. Recently, a number of solutions have been proposed including joint modelling and weighting [16, 17]; inverse proportional to attrition weighting is applied here.

We therefore aimed to distinguish the roles of childhood poverty and adult condition in explaining the trajectories of grip strength of Europeans aged 50 to 104 years. To tie the four strands together, three questions are raised. Do those with a poor childhood enter later life with a weaker grip and remain so throughout? Are their grip strength trajectories also steeper? Lastly, does good condition in adulthood render negligible any disadvantage identified earlier?

In answering these questions, this report contributes three ideas to the literature. The arm of childhood condition is long and strong in predicting grip strength much later in life. Childhood condition can be recovered retrospectively and should be considered when explaining health outcomes of people above 50. Epigenetic changes imprinted by poverty early in life may lie behind this long and strong arm of childhood condition.

## Materials and methods

The Survey of Health, Ageing, and Retirement in Europe (SHARE) is an ongoing longitudinal study of ageing in 20 countries so far [18]. Our use of this anonymised secondary research data has been approved for exemption by the ethical board of the University of Manchester.

As [6] we studied 11 countries, repeatedly surveyed and grouped into two regions: northern-continental (Austria, Denmark, France, Germany, the Netherlands, Sweden, Switzerland) and southern (Greece, Italy, and Spain). We used all waves (2004-2013) matched with the life course survey in 2008 following [5]. The matched sample differed from the rest in the following ways: the participants are older (67.0 vs 66.0 year, *t* = 17.5, *p* < 0.001) and somewhat weaker (33.5 vs 34.2 kg, *t* = 12.1, *p* < 0.001). There is a higher proportion of women to men in the analytic sample than in the excluded sample (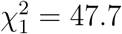, *p* < 0.001).

The outcome variable is objectively measured as the maximum grip strength of the dominant hand obtained using a dynamometer (Smedley, S Dynamometer, TTM, Tokyo, 100 kg) [6]. In contrast, childhood condition as the key exposure was retrospectively obtained. The condition concerned situation at ten years of age i.e. some four to nine decades in the past, indicating lack of the following: indoor toilet, hot and cold running water, central heating, fixed bath; plus overcrowding (more people than bedrooms) as well as number of books in the house, following [8].

It is tempting to use the information unmodified, but this should be resisted. A latent construct solution to obtaining poverty status when its indicators were inaccurate has been proposed [19]; a particular application has been fruitfully used on ELSA [8] as well as on the China Health and Retirement Longitudinal Study [20], sister studies of SHARE. Following this we built using latent class analysis a childhood poverty status giving poor versus non-poor class based on the indicators above. Beyond dealing with measurement error, this latent construct approach offers substantive advantages that we shall revisit in the discussion.

The literature on longitudinal ageing studies is keenly aware that participants tend to attrite non-randomly, hence a number of approaches have been proposed including pattern mixture [5], joint model [16, 21], multilevel multiple imputation [22, 23], and weighting [17, 24–26]. We joined the last stream to apply inverse proportional to attrition weighting. Specifically following [26] in their study of cognition in Atherosclerosis Risk in Community study, the attrition model includes age, sex, smoking, cognition, education, hypertension, cardiovascular disease, diabetes, and retirement status; stabilised weights were then computed with a base model including age, sex, and education.

The trajectories are derived using mixed model, also known as latent growth or random coefficients model, which has been used for this sample [5]. We included random intercepts only because there were no meaningful variations in the random age slopes nor extensive discussion of this in the literature [5, 7], retaining the virtue of parsimony [27]. Instead of positing that, ceteris paribus, the trajectories change randomly as age unfolds, we posited that they change systematically i.e. the childhood poor have a steeper decline.

We explored new factors unexamined in previous work on longitudinal trajectories of grip strength in SHARE. Social inequality in morbidities in later life is well documented, and this suggests inclusion of markers of socioeconomic position and marital status. Log of household income with purchasing power parity exchange rate, education (ISCED three levels: less than high school as reference, high school, and college or higher), occupation (ISCO three levels: elementary as reference, managerial or professional, and others), and marital status (fourfold: never married as reference, married or in partnership, separated or divorced, widowed). We included two markers of disadvantaged adult condition: following [5], adult occupational position (elementary occupation or not), and following [8], adult illness period.

Poverty class as derived above is one of the covariates. Because this is a derived latent class instead of an observed variable, adjustment to standard errors was made following a new method proposed by Vermunt and colleagues [28–30].

To answer the research questions, we built four models separately for men and women following [5–7]. The level model showed childhood poverty association with levels of grip strength, the slope model additionally showed association with the slope of annual decline by interacting poverty with age, while the alternative adult model showed, instead of the interaction term, additional adult condition associations. Lastly, the complete model includes them all. Modelling is done in Latent GOLD 5.1 [31] with model fit judged using Bayes-Schwarz information criterion of the smallest being best.

## Results

Women made up the majority of the sample (37,756, 55.6%) which has the average age of 66.9 years (standard deviation [SD]: 9.7 years). Women have weaker maximum grip: 26.0 kg (SD: 7.1 kg) compared to men: 42.6 kg (SD: 10.4 kg). The average of the maximum grip strength among northern-continental Europeans is (34.8 kg, SD: 12.0) while among southern Europeans is (31.2 kg, SD: 11.7 kg). The sample is further summarised in Table 1.

**Table 1.** Description of the analytic sample (SHARE 2002-2013)

| Variable | Women |  | Men |  |
| --- | --- | --- | --- | --- |
|  | Mean or *N* | Std. dev. or % | Mean or *N* | Std. dev. or % |
| Grip strength | 25.996 | 7.059 | 42.625 | 10.375 |
| Age | 66.836 | 10.029 | 67.042 | 9.338 |
| Height | 162.218 | 6.611 | 173.684 | 7.465 |
| Married | 24,361 | 64.52 | 24,511 | 81.22 |
| Single | 2,112 | 5.59 | 1,827 | 6.05 |
| Sep/divorced | 3,252 | 8.61 | 1,949 | 6.46 |
| Widowed | 8,031 | 21.27 | 1,892 | 6.27 |
| Elementary occ. | 996 | 2.64 | 539 | 1.79 |
| Intermediate | 36,068 | 95.53 | 28,574 | 94.68 |
| Professional | 692 | 1.83 | 1,066 | 3.53 |
| Primary or less | 14,185 | 37.57 | 9,284 | 30.76 |
| High School | 17,031 | 45.11 | 14,155 | 46.90 |
| College | 6,540 | 17.32 | 6,740 | 22.33 |
| Household income | 21,820 | 60,656 | 25,629 | 47,118 |
| Adult illness: none | 30,181 | 80.28 | 24,839 | 82.52 |
| One | 5,052 | 13.44 | 3,849 | 12.79 |
| Two | 1,022 | 2.72 | 734 | 2.44 |
| Three | 378 | 1.01 | 198 | 0.66 |
| More than three | 577 | 1.53 | 252 | 0.84 |
| Most adulthood | 384 | 1.02 | 230 | 0.76 |
| Adult elementary occ.: no | 27,644 | 73.22 | 21,060 | 69.78 |
| Yes | 10,112 | 26.78 | 9,119 | 30.22 |
| North | 23,259 | 61.60 | 18,525 | 61.38 |
| South | 14,497 | 38.40 | 11,654 | 38.62 |
| Lack facility: none | 6,580 | 17.43 | 5,466 | 18.11 |
| 1 | 3,999 | 10.59 | 3,012 | 9.98 |
| 2 | 3,475 | 9.20 | 2,708 | 8.97 |
| 3 | 5,691 | 15.07 | 4,560 | 15.11 |
| 4 | 7,786 | 20.62 | 6,270 | 20.78 |
| Lack all five | 10,225 | 27.08 | 8,163 | 27.05 |
| Num. books: none/very few | 17,103 | 45.78 | 13,794 | 46.13 |
| One shelf | 7,910 | 21.17 | 6,230 | 20.83 |
| One bookcase | 7,419 | 19.86 | 5,994 | 20.04 |
| Two bookcases | 2,483 | 6.65 | 1,870 | 6.25 |
| Three or more | 2,442 | 6.54 | 2,016 | 6.74 |
| Over-crowded: no | 10,103 | 26.76 | 8,312 | 27.54 |
| Yes | 27,653 | 73.24 | 21,867 | 72.46 |

The latent class analysis of childhood poverty revealed that 45.9% of the participants had a poor childhood at age ten (Table 2). The indicators of childhood condition showed plausible loadings. For instance, lacking more facilities is positively loaded on being a poor child while having more books is negatively loaded on being a poor child.

**Table 2.** Latent classes of poor and non-poor childhood.

| Indicator | Non-poor | Poor |
| --- | --- | --- |
| Size | 54.1% | 45.9% |
| Over-crowded |  |  |
| No | 0.4384 | 0.0597 |
| Yes | 0.5616 | 0.9403 |
| Lack facility |  |  |
| None | 0.2418 | 0.0116 |
| 1 | 0.1782 | 0.0228 |
| 2 | 0.1709 | 0.0567 |
| 3 | 0.1686 | 0.1398 |
| 4 | 0.1334 | 0.2673 |
| Lack all five | 0.1071 | 0.5019 |
| Number of books |  |  |
| None/very few | 0.2156 | 0.7091 |
| One shelf | 0.2377 | 0.2104 |
| One bookcase | 0.3245 | 0.0729 |
| Two bookcases | 0.1093 | 0.0061 |
| Three or more | 0.1129 | 0.0016 |

As women are found to have lower levels of grip strength than men, we presented their grip strength trajectories separately. The best model based on BIC is the adult model for both. Information criteria and key coefficients for both models are put together in Table 3 and discussed each in turn, putting complete information criteria (Supplement Table 4) and complete coefficients (Supplement Table 5) in the Supplement.

**Table 3.** Mixed models of trajectories of grip strength (adjusting for household income, occupation, education, and marital status); SE: standard error. Source: SHARE 2004-2013.

| Covariate | Women |  |  | Men |  |  |
| --- | --- | --- | --- | --- | --- | --- |
|  | coef | SE | *p* | coef | SE | *p* |
| Intercept | 28.1459 | 3.7292 | < 0.001 | 53.7396 | 3.3773 | < 0.001 |
| Age | -0.3028 | 0.0100 | < 0.001 | -0.4184 | 0.0112 | < 0.001 |
| Height | 0.0926 | 0.0212 | < 0.001 | 0.1303 | 0.0188 | < 0.001 |
| Southern Europe | -2.7182 | 0.2662 | < 0.001 | -2.5454 | 0.2862 | < 0.001 |
| Adult illness | -0.3664 | 0.0780 | < 0.001 | -0.6170 | 0.1445 | < 0.001 |
| Adult elementary occupation | -0.3857 | 0.0934 | < 0.001 | -0.7460 | 0.0801 | < 0.001 |
| Poor childhood | -1.1932 | 0.2370 | < 0.001 | -2.1917 | 0.2189 | < 0.001 |
| BIC | 359347 |  |  | 1990012 |  |  |

As has been widely documented, men have stronger grip (higher intercepts) but have steeper annual decline (418 versus 303 gram). Both slopes are significant (*p* < 0.001) and gentler than the estimates for the Danes [7]. Individual height and geographic region are also significant, in accordance with the literature [6]. Adult illness, a marker of adult condition, inversely associates with grip strength throughout; and so is being employed as elementary worker, which accords with [5]. Despite the strong effects of adult condition, the childhood-poor Europeans still have weaker grip in their later lives: men by 2.19 kg and women by 1.19 kg. In summary, the coefficients are plotted in Figure 1 to help in making comparison, and the trajectories of predicted grip strength for men and women who were childhood-poor and otherwise are drawn in Figure 2.

**Figure 1.**
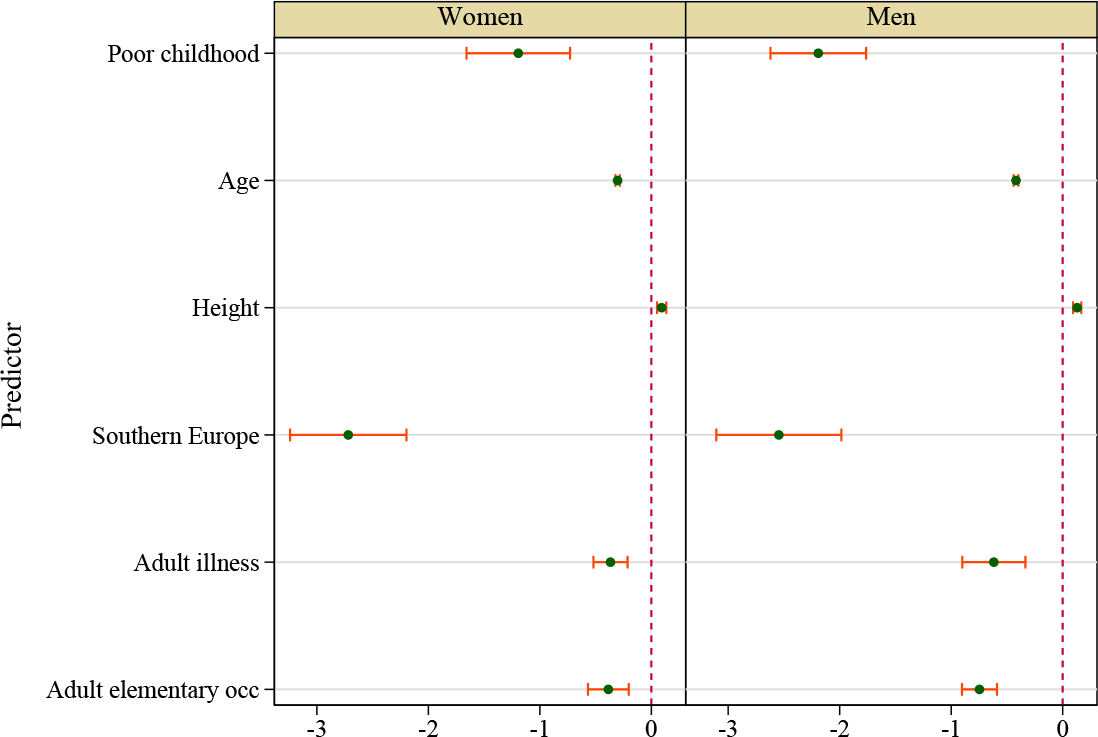
Plots of key coefficients for women (left pane) and men (right pane).

**Figure 2.**
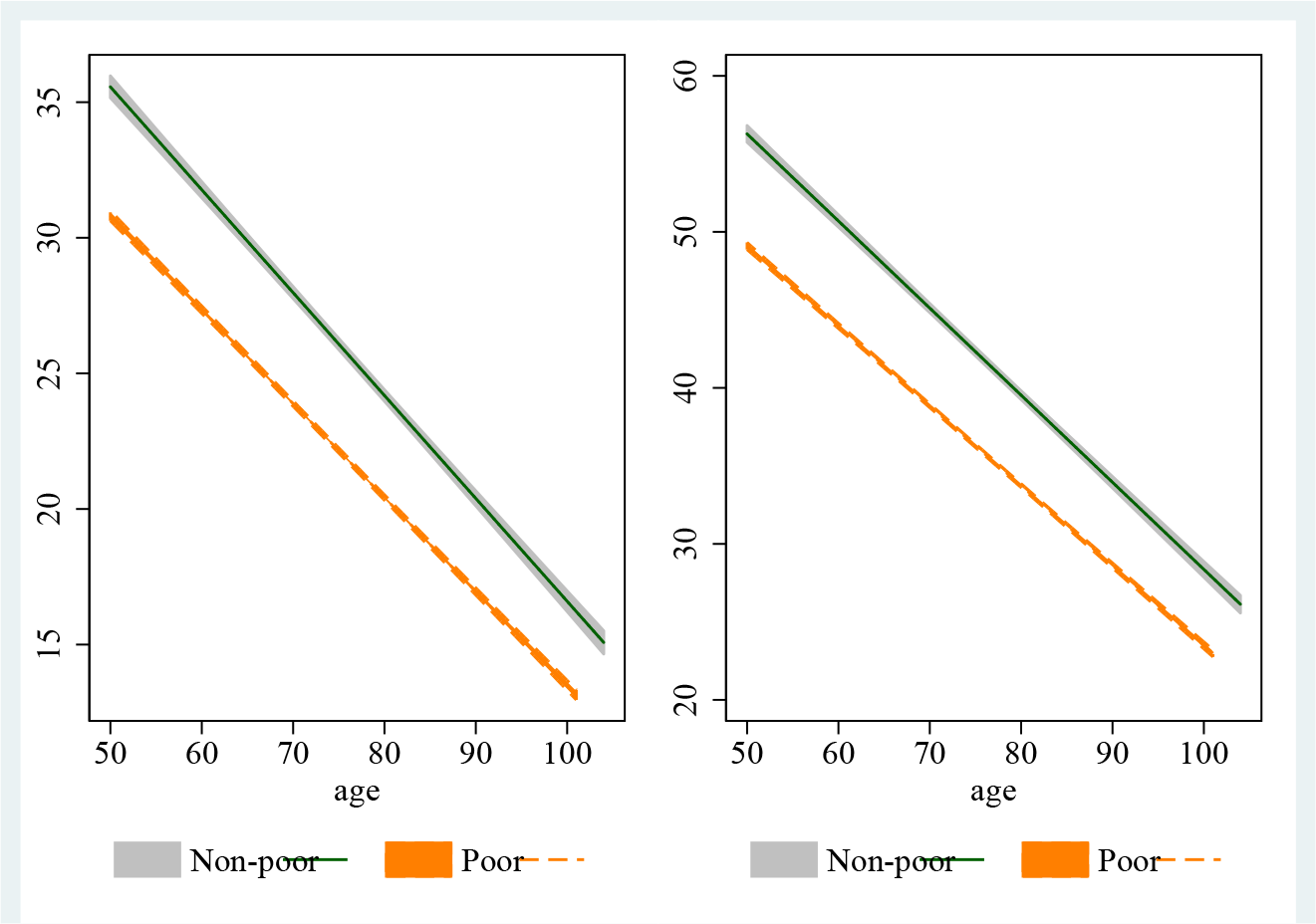
Predicted trajectories of grip strength, distinguished by childhood poverty status for women (left pane) and men (right pane).

## Discussion

Maintaining higher levels of grip strength is key, since it is a core component to avoid frailty and sarcopenia and ensure healthy ageing and wellbeing of older people. Here is the first evidence that being poor in life’s first decade goes with weaker grip in life’s last five decades.

Beyond covering a more extended age group than recent studies [5, 8], our study confirmed that adult condition (elementary occupation or ill health in adulthood) is associated with a weaker grip. These results are robust to inaccuracies in the measurement of childhood condition and to the attrition so common in longitudinal ageing studies. The results on the associations of childhood and adulthood conditions are strengthened because other factors have been accounted for including household income, education, occupational class, and marital status [32, 33]. In short, excepting the question about a steeper decline, the results supplied affirmative answers to all our questions: both childhood poverty and adulthood disadvantage go hand in hand with a weaker grip in later life.

Such long range results can be underpinned by a biosocial mechanism, especially with chronic inflammation playing a major role [34]. Older age is often marked by chronic or low grade inflammation which can impair muscle function. In turn, inflammation itself can be upregulated as a result of childhood adversity. The mechanism therefore has two major steps: [i] childhood adversity to lifetime inflammation, and [ii] inflammation disrupting myogenic processes of regeneration and functioning. We take each in turn.

Epigenetic literature has been accumulating evidence using animal models, such as mice, rats, and macaques, to examine whether early life adversity imprints epigenetic changes (DNA methylation and histone modification) to otherwise similar genotypes, resulting in different phenotypic response [34, 35]. Childhood poverty, pointing to broader early life adversity, entails more than just material lack but includes social deprivation when parents’ nurturing is compromised due to their time being absorbed in providing for household members and making ends meet.

Therefore animal models capable of reflecting some of the complexity of material and social deprivation are uniquely revealing, especially macaques studies. They have been used in a randomised design (of parental caring of frequent versus infrequent licking or grooming) to study the causal effect of early life deprivation on DNA methylation [36, 37]. The study found stable and organised epigenetic changes, involving genes in the pathways of the immune system and the hypothalamic pitutitary adrenal (HPA) axis responsible for responding to stress. The peripheral immune system interacts with the HPA axis and has a role in brain function; evidence consistent with this interaction has been shown in this sample in our previous work [38].

A key gene for regulating the HPA axis function, the glucocorticoid receptor (*NR*3*C*1), is activated in the hypothalamus in response to stress and releases glucocorticoid. Glucocorticoid receptor is differentially expressed according to the experience of social deprivation, by epigenetic programming through histone acetylation and DNA methylation of the exon **1_F_**. This epigenetic programming differentiates similar DNA sequences phenotypically, resulting in blunted feedback by glucocorticoid and heightened stress response and demodulated immune system response, a pattern that is stable throughout the life course. The bidirectional interaction between the HPA axis and the immune system facilitates the imprinting of childhood adversity through epigenetic changes. This can lead to chronic inflammation that is stable through later life as reflected in higher levels of circulating tumour necrosis factor-*α* (TNF-*α*).

By discussing the role of inflammatory cytokines such as TNF-*α*, the literature on muscle regeneration and muscle function has provided evidence to complete the mechanism. Inflammation is known to impair both muscle regeneration and muscle functioning. In normal activities of daily living which involve muscle exertion, some minute damage to muscle tissue may occur [39]. In these circumstances, the pluripotent myosatellite cells respond by proliferating and differentiating to form muscle fibres and cover the damaged tissue. But circulating inflammatory cytokines such as TNF-*α* have been shown to impair this process of regeneration in two ways: apoptosis of myoblasts [40] and inhibition of the differentiation stage, leaving proliferated cells unable to differentiate and replace the damaged tissue [41]. Beyond impairing the myogenesis process in common minute damage, inflammation also impairs functioning by reducing the power of the single permeable fibre [42]. So in mice, TNF-*α* rapidly reduces the force generating capacity or specific tension of muscle fibres independent of loss of muscle volume [43].

In short, inflammation impairs muscle functioning in older people at least along three points: it encourages myosatellite cell deaths [44], it interrupts the step of differentiation into myonuclei and muscle fibres; lastly, even if muscle fibres were successfully regenerated, inflammation reduces the febrile tensile strength. Childhood poverty, through upregulating inflammation, impairs grip strength in later life.

This study has a number of weaknessess. First, by matching only individuals with childhood information with those with longitudinal observations, inevitably some unmatched observations were set aside. It is impossible to measure the direction of possible bias this might entail.

Second, although epigenetic changes are posited to be the mechanism, there is no direct evidence of the extent of DNA methylation in the sample. This is a potentially rectifiable weakness. Despite these weaknesses, this study has some strengths. First, the sample is designed to represent the countries and not only some clinical groups or cities, hence facilitating generalisation. Finally, this study is also the first to link broad childhood condition (subject to recall error) with later life trajectories (subject to attrition), reinforcing sustained links across the life course.

As alluded to above, besides uncovering the strong results on childhood poverty, the method with which childhood poverty is constructed i.e. as a latent class of poverty, holds potential to advance research work on the life course and health. It is useful to call to mind that childhood information, such as a lack of the five facilities above, can be used alternatively as (i) indicators of a latent factor in factor analysis or (ii) five additional covariates. Now the use of a latent class of poverty facilitates discussion, for instance when presenting whether the childhood poor (compared to the non-poor) show better health outcomes in later life. On the other hand, with the latent factor we have to compare those on one standard deviation away from the mean against those on the mean of the latent factor. This is hardly intuitive. Or with five additional covariates, we are led to scrutinise each effect which makes discussion potentially unwieldy.

Second, a latent class is also easier to use when testing a hypothesis of a steeper decline; it simply needs an interaction term (of poverty class and age). The interpretation will be similarly intuitive: the childhood poor declined more steeply by a certain kg (the coefficient) per year if the interaction term was found significantly negative. On the other hand, although with the latent factor a similar interaction can be used, there remains the attendant difficulty of interpretation. Or with five additional covariates, we are required to use five interaction terms. Depending on the choice, interpretation may be hindered.

Most important, a latent class enables cross-country comparison. It is conceivable that the long arm of childhood condition hypothesis may be tested in other countries, for example in equatorial developing countries where lack of running hot water or central heating may not hold similar salience. Without these indicators, nevertheless, poverty class can still be constructed with this method and the hypothesis tested. In this way, latent class of childhood poverty based on retrospective information should always be considered in life course and ageing investigation anywhere in the world.

In conclusion, although childhood and adulthood conditions last a lifetime [12], there is a potential role for interventions in adulthood both in the labour market and the health sector. On the basis of evidence uncovered here, childhood could be a critical period to stave costly long term care. It is never too early to invest in later life.

## Acknowledgments

This work was supported by grants from the Medical Research Council and Economic & Social Research Council (No. G1001375/1), the National Institute of Health Research (No. ES/L001772/1) and the European Union’s Horizon 2020 Research and Innovation Programme (No. 668648) to Gindo Tampubolon.

This paper used data from SHARE Waves 1, 2, 3 (SHARELIFE), 4 and 5 (DOIs: 10.6103/SHARE.w1.500, 10.6103/SHARE.w2.500, 10.6103/SHARE.w3.500, 10.6103/SHARE.w4.500, 10.6103/SHARE.w5.500), see Börsch-Supan et al. (2013) for methodological details. The SHARE data collection has been primarily funded by the European Commission through the FP5 (QLK6-CT-2001-00360), FP6 (SHARE-I3: RII-CT-2006-062193, COMPARE: CIT5-CT-2005-028857, SHARELIFE: CIT4-CT-2006-028812) and FP7 (SHARE-PREP: No. 211909, SHARE-LEAP: No. 227822, SHARE M4: No. 261982). Additional funding from the German Ministry of Education and Research, the U.S. National Institute on Aging (U01_AG09740-13S2, P01_AG005842, P01_AG08291, P30_AG12815, R21_AG025169, Y1-AG-4553-01, IAG_BSR06-11, OGHA_04-064) and from various national funding sources is gratefully acknowledged (see www.share-project.org)

## Supplement Table 4 and 5

**Table 4.**
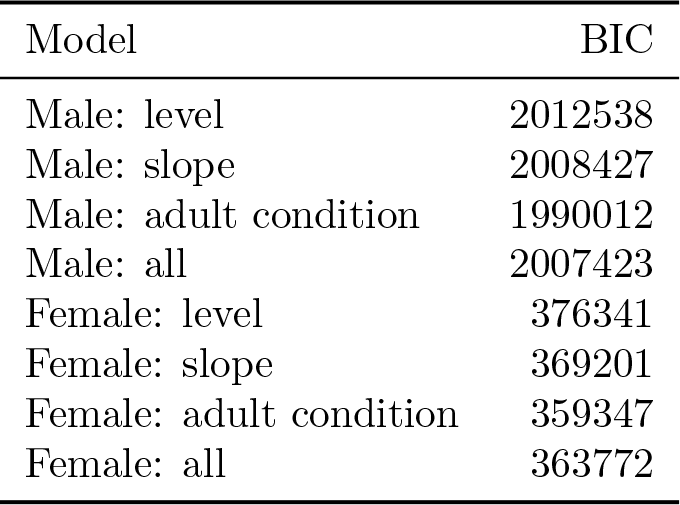
Supplement Table 4 Model comparisons.

| Model | BIC |
| --- | --- |
| Male: level | 2012538 |
| Male: slope | 2008427 |
| Male: adult condition | 1990012 |
| Male: all | 2007423 |
| Female: level | 376341 |
| Female: slope | 369201 |
| Female: adult condition | 359347 |
| Female: all | 363772 |

**Table 5.** Supplement Table 5 Models with adult condition for women (left pane) and men (right pane)

| Covariate | Women |  |  | Men |  |  |
| --- | --- | --- | --- | --- | --- | --- |
|  | coef | SE | *p* | coef | SE | *p* |
| Intercept | 28.1459 | 3.7292 | < 0.001 | 53.7396 | 3.3773 | < 0.001 |
| Poor childhood | -1.1932 | 0.2370 | < 0.001 | -2.1917 | 0.2189 | < 0.001 |
| Age | -0.3028 | 0.0100 | < 0.001 | -0.4184 | 0.0112 | < 0.001 |
| Log household income | 0.1375 | 0.0026 | < 0.001 | 0.1633 | 0.0148 | < 0.001 |
| High school | 2.2142 | 0.2716 | < 0.001 | 1.9144 | 0.2723 | < 0.001 |
| College | 5.5721 | 0.3256 | < 0.001 | 10.2978 | 0.5578 | < 0.001 |
| Intermediate | 1.7347 | 0.1905 | < 0.001 | -1.8704 | 0.3893 | < 0.001 |
| Managerial | 3.7451 | 0.3295 | < 0.001 | 0.2778 | 0.5475 | 0.61 |
| Married | 1.9782 | 0.3128 | < 0.001 | -2.1077 | 0.4301 | < 0.001 |
| Sep/divorced | 4.7904 | 0.7347 | < 0.001 | 4.3090 | 0.3546 | < 0.001 |
| Widowed | 2.0289 | 0.3310 | < 0.001 | -3.5030 | 0.5218 | < 0.001 |
| Height | 0.0926 | 0.0212 | < 0.001 | 0.1303 | 0.0188 | < 0.001 |
| Southern Europe | -2.7182 | 0.2662 | < 0.001 | -2.5454 | 0.2862 | < 0.001 |
| Adult illness period | -0.3664 | 0.0780 | < 0.001 | -0.6170 | 0.1445 | < 0.001 |
| Adult elementary work | -0.3857 | 0.0934 | < 0.001 | -0.7460 | 0.0801 | < 0.001 |
| $\sigma^2$ | 4.1348 | 0.0805 | < 0.001 | 4.1061 | 0.0940 | < 0.001 |

